# Identification of a major QTL and associated marker for high arabinoxylan fibre in white wheat flour

**DOI:** 10.1101/705343

**Authors:** Alison Lovegrove, Luzie U. Wingen, Amy Plummer, Abigail Wood, Diana Passmore, Ondrej Kosik, Jackie Freeman, Rowan A.C. Mitchell, Mehmet Ulker, Karolina Tremmel-Bede, Marianna Rakszegi, Zoltán Bedő, Marie-Reine Petterant, Gilles Charmet, Michelle Leverington Waite, Simon Orford, Amanda Burridge, Till Pellny, Peter R Shewry, Simon Griffiths

**Affiliations:** Rothamsted Research, West Common, Herts AL5 2JQ; John Innes Centre, Norwich Research Park, Colney Lane, Norwich NR4 7UH; Bardakçi Mahallesi, Yüzüncü Yil Üniversitesi Kampüsü, 65090 Tuşba/Van, Turkey; Centre for Agricultural Research, Hungarian Academy of Sceinces, Martonvásár, Hungary; INRA, 5 Chemin de Beaulieu, 63000 Clermont-Ferrand, France; Life Sciences, University of Bristol, 24 Tyndall Avenue, Bristol, BS8 1TQ

**Keywords:** Wheat, white flour, dietary fibre, arabinoxylan, relative viscosity, QTL

## Abstract

Dietary fibre (DF) has multiple health benefits, and wheat products are major sources of DF for human health. However, DF is depleted in white flour, which is most widely consumed, compared to wholegrain. The major type of DF in white wheat flour is the cell wall polysaccharide arabinoxylan (AX). Previous studies have identified the Chinese wheat cultivar Yumai 34 as having unusually high contents of AX in both water-soluble and insoluble forms. We have therefore used populations generated from crosses between Yumai 34 and four other wheat cultivars, three with average contents of AX (Ukrainka, Altigo and Claire) and one also having unusually high AX (Valoris), in order to map QTLs for soluble AX (determined as relative viscosity) of aqueous extracts of wholemeal flours) and total AX (determined by enzyme fingerprinting of white flour). A number of QTL were mapped, but most were only detected in one or two crosses. However, all four crosses showed strong QTLs for high RV/total AX on chromosome 1B, with Yumai 34 being the increasing parent, and a KASP marker for the high AX Yumai 34 allele was validated by analysis of high AX lines derived from Yumai 34 but selected by biochemical analysis. A QTL for RV was mapped on chromosome 6B in Yumai 34 × Valoris, with Valoris being the increasing allele, which was consistent with the observation of transgressive segregation for this trait. The data indicate that breeding can be used to develop wheat with high AX fibre in white flour.

## Introduction

Dietary fibre (DF) is essential for human health, with cereals providing about 40% of the total fibre intake in Western European countries such as the UK (Bates et al., 2014) and Finland (Helldan et al., 2012). Dietary fibre (DF), and wholegrain cereal fibre in particular, has been shown to have a number of health benefits, including lowering blood pressure and serum cholesterol, improving insulin sensitivity and reducing the incidence of certain types of cancer, notably bowel and breast cancers (Cade et al, 2007; Wood, 2007; Anderson et al, 2009; Wolever et al, 2010; Aune et al, 2011; 2016; Cooper et al., 2015; Ye et al., 2012; Hajishafiee et al., 2016; Reynolds et al., 2018)

The mechanisms are still incompletely understood, but are considered to include increasing faecal bulk and reducing intestinal transit time, binding cholesterol and carcinogens, reducing the rate of digestion and glucose release in the small intestine and fermentation to beneficial short chain fatty acids in the colon. DF also occurs in soluble and insoluble forms, which are considered to differ in some respects in their health benefits, with insoluble fibre being more slowly fermented and contributing particularly to binding cholesterol and carcinogens and increasing faecal bulk. However, despite these established health benefits, the intake of DF in most countries falls far below the recommended levels. For example, the daily intake in the UK is currently 17.2g/day for women and 20.1 g/day for men, compared with a recommended intake of 30g/day (https://www.nutrition.org.uk/nutritionscience/nutrients-food-and-ingredients/dietary-fibre.html).

Although wholegrain wheat is relatively rich in fibre, containing about 11 to 15% dry weight, (Andersson et al., 2012), most wheat products are made from white flour (derived from the starchy endosperm) (Steer et al., 2008) which contains only 2-3% fibre (Gebruers et al, 2008). Furthermore, increased consumption of highly refined cereal products (including bread and other products from white flour) is occurring in countries undergoing urbanisation and industrialisation, which is considered to contribute to increases in obesity and chronic diseases in these countries (Mattei et al, 2015).

The major components of the DF fraction of wheat flour are cell wall polysaccharides, principally arabinoxylan (AX) and (1→3,1→4)-β-D-glucan (β-glucan), which account for about 70% and 20% of the total, respectively, with about 2% cellulose ((1→4)-β-D-glucan) and 7% glucomannan (Mares and Stone, 1973). However, the content of AX also varies between different genotypes of wheat. For example, 2-fold variation in the content of total (TOT)-AX and 4.7-fold variation in water-extractable (WE)-AX was reported in white flour of 150 wheat genotypes grown together on a single site (Gebruers et al., 2008), and 2.9-fold variation in WE-AX and 1.7-fold variation in water-unextractable (WU)-AX in 20 wheat cultivars (Ortiz-Ordaz and Saulnier, 2005). Furthermore, a high proportion of the variation in the AX content of wholemeal and white flours of wheat is heritable, and hence accessible for exploitation by breeders (Martinant et al., 1999; Shewry et al., 2010). However, the exploitation of this variation to develop improved wheats has been limited by the lack of tools for selection, with biochemical analyses being slow and costly and a lack of molecular markers for selection.

A number of studies of the genetic control of AX content have been reported using genetic analysis, with most analysing wholemeal flour by either direct determination of AX (by monosaccharide analysis or colorimetric determination) or the relative viscosity of aqueous extracts as a proxy (Martinant et al. 1998; Perretant et al. 2000; Laperche et al. 2007; Quraishi et al. 2009; Charmet et al. 2009; Nyugen et al; Yang et al., 2015), for AX content. In addition, two association studies of AX in wholemeal tetraploid wheat (Marcotuli et al., 2015) and in white flour of bread wheat (Quraishi et al., 2011) have been reported. However, although these studies identified a number of QTL, these were not consistent across crosses and have not led to the identification of markers for breeding.

We have therefore determined the genetic control of AX content in white flour of wheat, by exploiting crosses with the high AX Chinese cultivar Yumai 34 (Gebruers et al, 2008). Analysis of crosses between this cultivar and three cultivars with normal levels of fibre identified a major QTL on chromosome 1B, while analysis of a cross with a second high AX genotype (Valoris) identified a second major QTL (on chromosome 6B). This allowed the identification of a linked marker for the 1B QTL which was validated by analysis of high AX lines developed from Yumai 34 using biochemical analysis for selection.

## Results

### Identification of Yumai 34 as a source of high arabinoxylan fibre in white flour

Analysis of 150 bread wheat cultivars in the EU Healthgrain project identified the Chinese wheat cultivar Yumai 34 as having the highest contents of both WE-AX and total AX in white flour, with 1.4% water-extractable AX (WE-AX) and 2.75% total AX (TOT-AX), compared with mean values for the 150 lines of 0.51% and 1.93%, respectively (Gebruers et al, 2008). The high proportion of WE-AX was especially notable, corresponding to over 50% of TOT-AX in Yumai 34 compared with a mean of 26% for the 150 lines.

Although these analyses were carried out on grain samples from single plots grown on the same site in 2005-6, further comparative analyses carried out on lines grown for over 10 years has confirmed that Yumai 34 always contains the highest, or among the highest, contents of both TOT and WE- AX fractions in flour (authors’ unpublished results). This is illustrated by Table 1, which compares the contents of AX fractions in white flour of four cultivars grown at Rothamsted in 2011/2012. Yumai 34 clearly has the highest contents of TOT-AX and WE-AX, with the latter being reflected in the high relative viscosity (RV) of aqueous extracts (as discussed below).

**Table 1.** Comparison of the contents of TOT-AX and WE-AX, the arabinose:xylose (A:X ratio*) of TOT-AX and the relative viscosity of aqueous extracts of white flour from four wheat cultivars grown in a field trial in 2011/2012.

| Cultivar | TOT-AX<br>(% dry wt) | WE-AX<br>(% dry wt) | A:X Ratio<br>of TOT-AX | Relative<br>viscosity<br>of<br>aqueous<br>extracts |
| --- | --- | --- | --- | --- |
| Yumai 34 | 2.89 | 1.34 | 0.44 | 2.77 |
| Manital | 1.77 | 0.69 | 0.52 | 1.70 |
| Hereward | 1.61 | 0.73 | 0.49 | 1.75 |
| Soissons | 1.62 | 0.58 | 0.53 | 1.62 |

\* Adjusted for AGP (Gebruers et al., 2009)

Arabinoxylan comprises a backbone of β-D-xylopyranosyl residues linked through (1→4) glycosidic linkages with some residues being substituted with α-L-arabinofuranosyl residues at either position 3 or positions 2 and 3. Monosaccharide analysis showed that the ratio of arabinose to xylose (A:X) in TOT-AX is lower for Yumai 34 than for other cultivars, indicating that the structure of AX also differs (Table 1). This difference was therefore investigated further using enzyme fingerprinting. Treatment of AX with a specific type of endoxylanase enzyme results in cleavage of the xylan backbone, releasing a mixture of xylose, xylobiose, xylotriose (comprising 1, 2 and 3 xylose units, respectively) and arabinoxylan-derived oligosaccharides (AXOS) comprising 4 to 7 xylose units, one or more of which may be mono- or disubstituted with arabinose. The proportions of these AXOS therefore provides a “fingerprint” which reflects differences in the extent and pattern of arabinose substitution. The application of this approach to AX from white flours of the four cultivars showed that Yumai 34 has a high proportion of a pentasaccharide with the structure XA3XX (xylose-xylose monosubstituted with arabinose-xylose-xylose), which comprised 26% of the total fragments compared with 15.17%, 16.14% and 17.21% in the other cultivars (Supplementary Table S1). This increase is associated with reduced proportions of oligosaccharides containing disubstituted arabinose residues, and accounts for the lower A:X ratio shown in Table 1.

### Development of populations for genetic analysis of AX

Four populations were generated from crosses between Yumai 34 and European cultivars: with the central European cultivar Ukrainka (96 F6 recombinant inbred lines (RILs)) (Y34Ukr), the UK biscuit-making cultivar Claire (95 RILs) (Y34Cl) and the French breadmaking cultivars Altigo (245 doubled haploid lines (DHL)) (Y34Alt) and Valoris (84 DHL) (Y34Val). Whereas Ukrainka, Altigo and Claire and were selected as parents because they had average contents of AX, previous studies have shown that Valoris has higher than average contents of both WE-AX and TOT-AX (0.8% and 2.2% compared with means of 0.51% and 1.93%, respectively, in the Healthgrain study) and it has therefore been used as a “high AX parent” in crosses with other low AX genotypes (Charmet et al., 2009).

### Mapping relative viscosity (RV) of aqueous extracts of wholemeal

Most previous genetic analyses of AX fibre in wheat have determined the RV of aqueous extracts of white or wholemeal flours as a proxy for arabinoxylan content (reviewed by Quraishi et al., 2010; Shewry, 2013). This is because this parameter is more readily determined than AX, eliminating the need for milling and biochemical analysis. It is considered to be valid because WE-AX is known to be the major contributor to the viscosity of aqueous extracts. However, the use of RV of wholemeal flour as a proxy for WE-AX in white flour has not been validated. We therefore compared the RV of aqueous extracts and the contents of WE-AX in wholemeal and white flours from 10 lines from the Y34Ukr cross (grown at Rothamsted in 2013/4). This showed good correlations between wholemeal RV and wholemeal WE-AX (determined as pentosans) (r=0.70), white flour RV (r=0.84) and white flour WE-AX (pentosans) (0.77) (Supplementary Figure S1)

The RV of aqueous extracts from single samples of wholemeal flours from the four populations was therefore determined. This showed clear transgressive segregation in the population derived from the cross with Valoris (which has high AX) (Figure 1), but not in the populations derived from crosses between Yumai 34 and the average AX lines: Altigo, (Figure 1), Claire and Ukrainka (Supplementary Figure S2).

**Figure 1.**
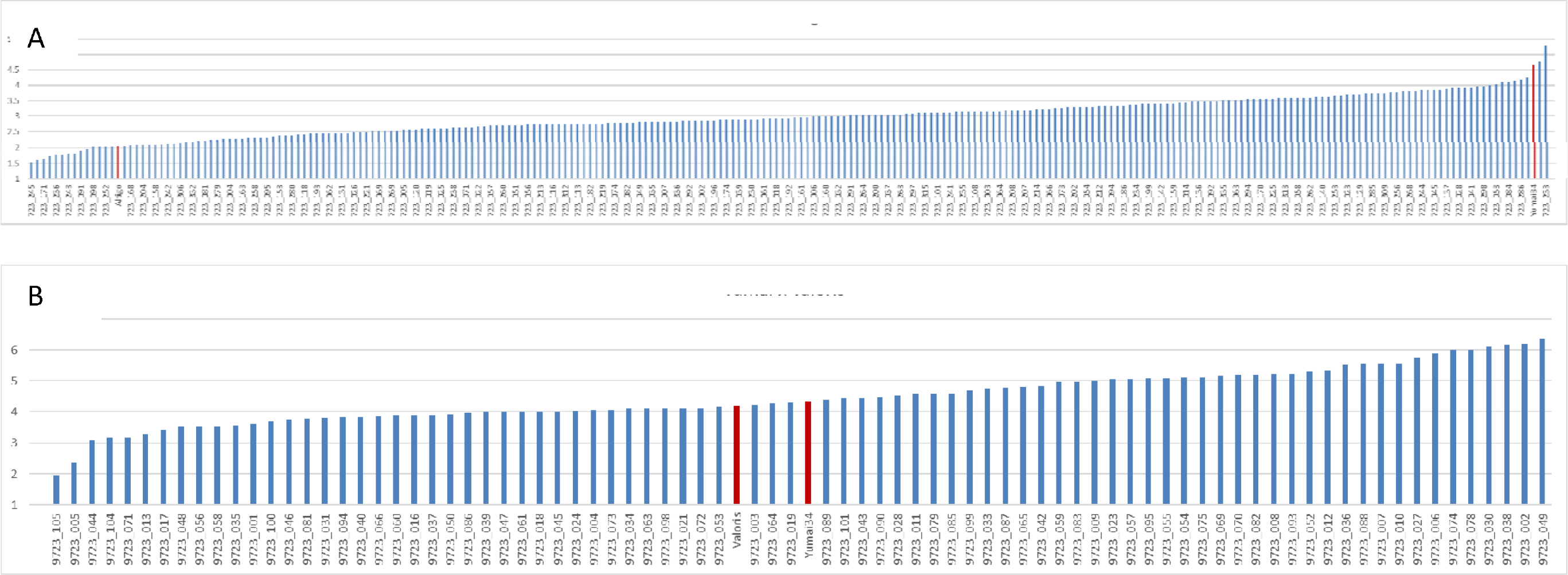
Relative viscosity of wholemeals flours from DH populations from the crosses Yumai 34 × Altigo (A) and Yumai 34 × Valoris (B), Parental lines are shown in red.

The QTL identified in these crosses are shown in Table 2. Seven RV QTL were identified on chromosomes 1AS, 1BL, 2BS, 2D, 3BL, 4DL, and 6BS: four in Y34Alt, 2 in Y34Cl, none in Y34Ukr and 3 in Y34Val. These QTL positions are projected onto the genome of Chinese Spring (Appels et al 2018) in Figure 2. The relative mapped positions of flanking and peak markers for 2D QTL may suggest rearrangement. Where the QTL 1 LOD confidence intervals substantially overlap (green line on figure) for the same trait in different populations we have made the assumption that the underlying effect is the same. The increasing alleles for these QTL come from Yumai 34 except for a QTL for RV on 1A in Y34Val and a QTL for RV on 6B in Y34Val.

**Figure 2.**
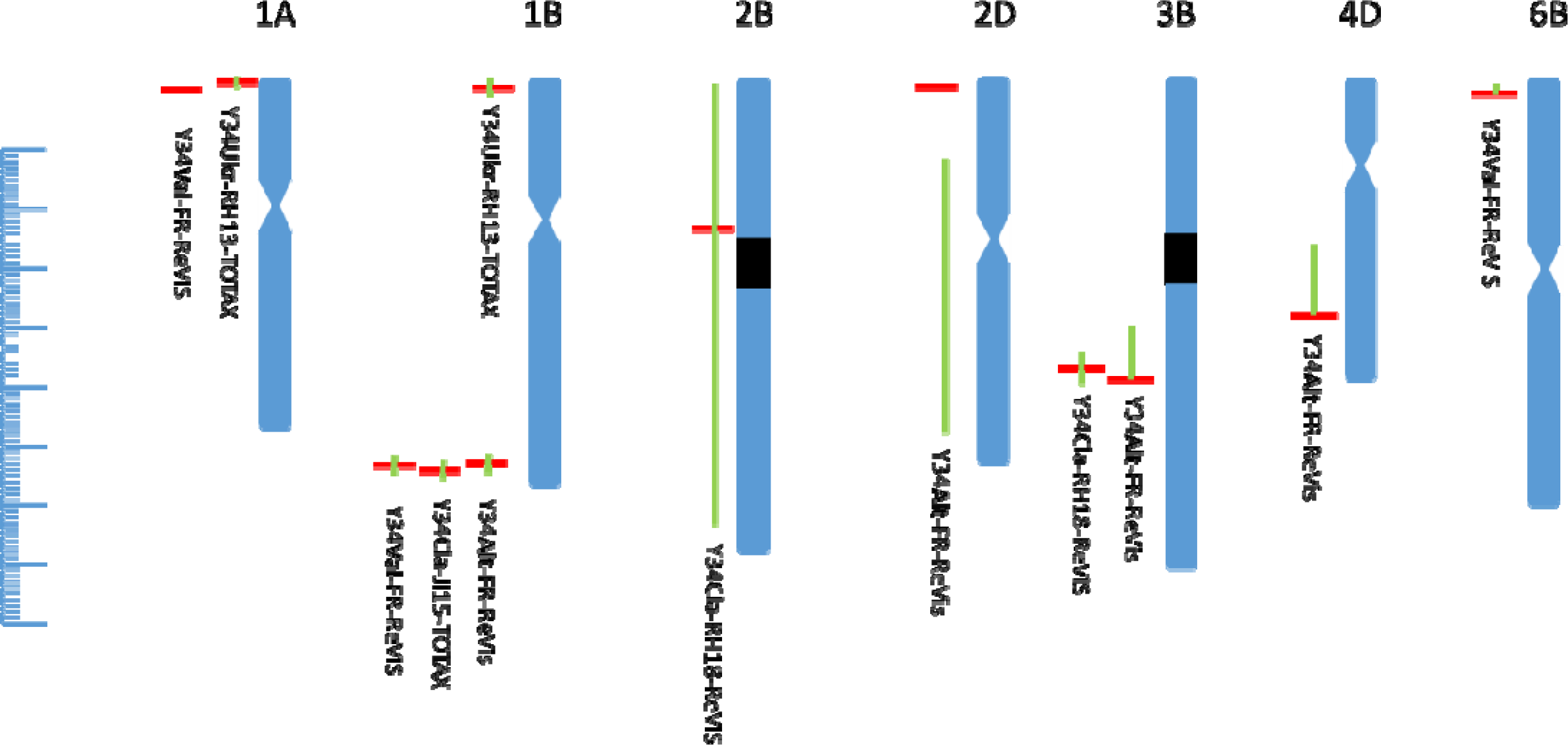
Chromosomal locations of QTLs for RV and TOT-AX mapped in the four crosses. Alignment of QTL peak marker (horizontal red bar) and flanking markers defining 1 LOD confidence interval (edges of green vertical bar) is to Chinese RefSeq v1.0. Each hash mark on the left-hand scale represents 1Mb. Approximate location of centromeres is shown as a hour glass when determined using chromosome arm survey sequence data (International Wheat Genome Sequencing Consortium, 2014) and as black blocks for 2B and 3B using density of annotated genes (Appels et al 2018). QTL are named as Population-environment-trait (see Table 2).

**Table 2.** QTLs above LOD threshold 3 for relative viscosity (RV) and TOT-AX in four crosses Y34; Yumai 34; Alt, Altigo; Cl, Claire; Ukr, Ukrainka; Val, Valoris. Chr is the chromosome on which QTL are identified env is the environment in which the QTL was detected (location and year): FR, Clermont Ferrand, JI, John Innes Centre, RH, Rothamsted. % var is the percentage of phenotypic variance explained, mean is the mean of the trait for the whole population, increasing allele indicates the direction of the allelic effect.

| Population | Chr | env | Trait | LOD | %var | add eff | mean | increasing Allele |
| --- | --- | --- | --- | --- | --- | --- | --- | --- |
| Y34Alt | 1B | FR2014 | RV | 12.6 | 16.3 | -0.53 | 3.0 | Y34 |
|  | 2D | FR2014 | RV | 3.2 | 3.7 | -0.26 | 3.0 | Y34 |
|  | 3B | FR2014 | RV | 6.1 | 7.4 | -0.3 | 3.0 | Y34 |
|  | 4D | FR2014 | RV | 3.5 | 4.1 | -0.27 | 3.0 | Y34 |
| Y34CI | 2B | RH2018 | RV | 3.1 | 11.4 | -0.038 | 1.8 | Y34 |
|  | 3B | RH2018 | RV | 5.9 | 23.9 | -0.013 | 1.8 | Y34 |
|  | 1B | JI2015 | TOT-AX | 3.2 | 14.1 | -3.8 | 74.4 | Y34 |
| Y34Ukr | 1A | RH2013 | TOT-AX | 3.2 | 13.0 | 2.1 | 95.1 | Ukr |
|  | 1B | RH2013 | TOT-AX | 5.1 | 21.6 | -3.0 | 95.1 | Y34 |
| Y34Val | 1A | FR2014 | RV | 3.8 | 10.3 | 0.62 | 4.5 | Val |
|  | 1B | FR2014 | RV | 7.8 | 24.2 | -0.99 | 4.5 | Y34 |
|  | 6B | FR2014 | RV | 4.4 | 12.3 | 0.85 | 4.5 | Val |

A major objective of this study was to understand the genetic basis of the high RV of Yumai 34 so QTL with increasing alleles from this variety, of large effect, and expressed in multiple populations are of particular interest. In this respect 1BL stands out at LOD 12.6 and 7.8 in the Y34Alt and Y34Val populations, respectively, accounting for 16.3% and 24.2% phenotypic variance. With additive effects of 0.53 and 0.99 RV units these QTL alone can deliver substitution effects of 1 to 2 RV units in populations with mean RVs of 3 and 4.5. A weaker RV effect also detected in two populations, Y34Alt and Y34Cl, is found on 3B. No other RV effects are detected in more than one population.

### Mapping TOT-AX determined by fingerprinting

Analyses of ten Y34Ukr lines (discussed above) showed that RV of aqueous extracts of wholemeal was a poor predictor of TOT-AX (calculated as the sum of the AXOS released) in white flour (r=0.38) (Supplementary Figure S1). We therefore determined the amount of TOT-AX by enzyme fingerprinting of 3 replicate samples of the Y34Ukr, Y34Cl and Y34Val populations.

Three TOT-AX QTL were identified in the Y34Ukr and Y34Cl populations. Both populations segregated for a TOT-AX QTL on 1B with Yumai 34 carrying the increasing allele. However, alignment to the wheat genome sequence (Fig 2) shows that the location on 1B are very different, sub-telomeric 1BL in Y34Cl and sub-telomeric 1BS in Y34Ukr. For the third TOT-AX QTL Ukrainka carries the increasing allele on 1AS.

### Coincidence of RV and TOT-AX QTL

Fig 2 shows how some of the QTL for the two traits mapped in the four populations co-locate. The Claire TOT-AX QTL on 1BL co-locates with the QTL for RV from Y34Alt and Y34Val, with Yumai 34 carrying the increasing allele in all cases. Two further co-locating QTL are not shown in Table 2 or Figure 2 because the LOD scores were below the cut-off. These were a QTL for TOT-AX in Y34Val (LOD 2.5) and a QTL for RV in Y34Ukr (LOD2.3). Interestingly, a second TOT-AX QTL from Y34Ukr is located on the opposite end of chromosome 1B (1BS).

QTLs for RV with Yumai 34 carrying the increasing allele were co-located on chromosome 3B in Y34Alt and Y34Cl and were also identified on chromosomes 2B (Y34Alt), 4D (Y34Alt) and 2B (Y34Cla).

Three QTL were identified in which Yumai 34 was not the increasing parent, for TOT-AX and RV from Y34Ukr and Y34Val, respectively, which co-located on chromosome 1A and for RV on chromosome 6B in Y34Val only.

### Identification and validation of markers for the high AX QTL from Yumai 34

The QTLs identified on 1BL for which Yumai 34 alleles increased TOT-AX and/or RV were prioritised for the development of KASP SNP assays which would allow validation of the QTL effect in independent populations and the selection of high AX Yumai 34 alleles in breeding programmes. The five Axiom markers nearest to the RV QTL peak of Y34Alt were therefore selected for KASP marker development. The SNP BA00789946 detected by AX-94524314 was converted into a robust KASP assay and used to genotype the Y34Alt population, giving exactly the same segregation pattern as the corresponding Axiom marker.

The marker was then validated using high fibre breeding lines developed by Tremmel-Bede et al. (2017) from crosses between Yumai 34 and conventional cultivars. These lines were selected by determining the contents of WE-AX and TOT-AX in white flour, showing increases in WE-AX of about 0.5% and in TOT-AX of about 1% compared to the conventional parents. Of 11 high fibre lines selected from a cross between Yumai 34 and the Ukrainian winter wheat cultivar Ukrainka, 9 carried the Yumai 34 allele of the BA00789946 KASP marker. The alleles in these lines are shown together with the contents of WE-AX in white flour in Figure 3. A second set of selections from the same study was genotyped from a cross between Yumai 34 and the German wheat cultivar Lupus. Although no polymorphism was detected using DNA from available accessions of Yumai 34 and Lupus, with both carrying the Yumai34 allele of BA00789946, the nine high-fibre progeny did segregate with eight carrying the Yumai 34 allele and one the alternative allele.

**Figure 3.**
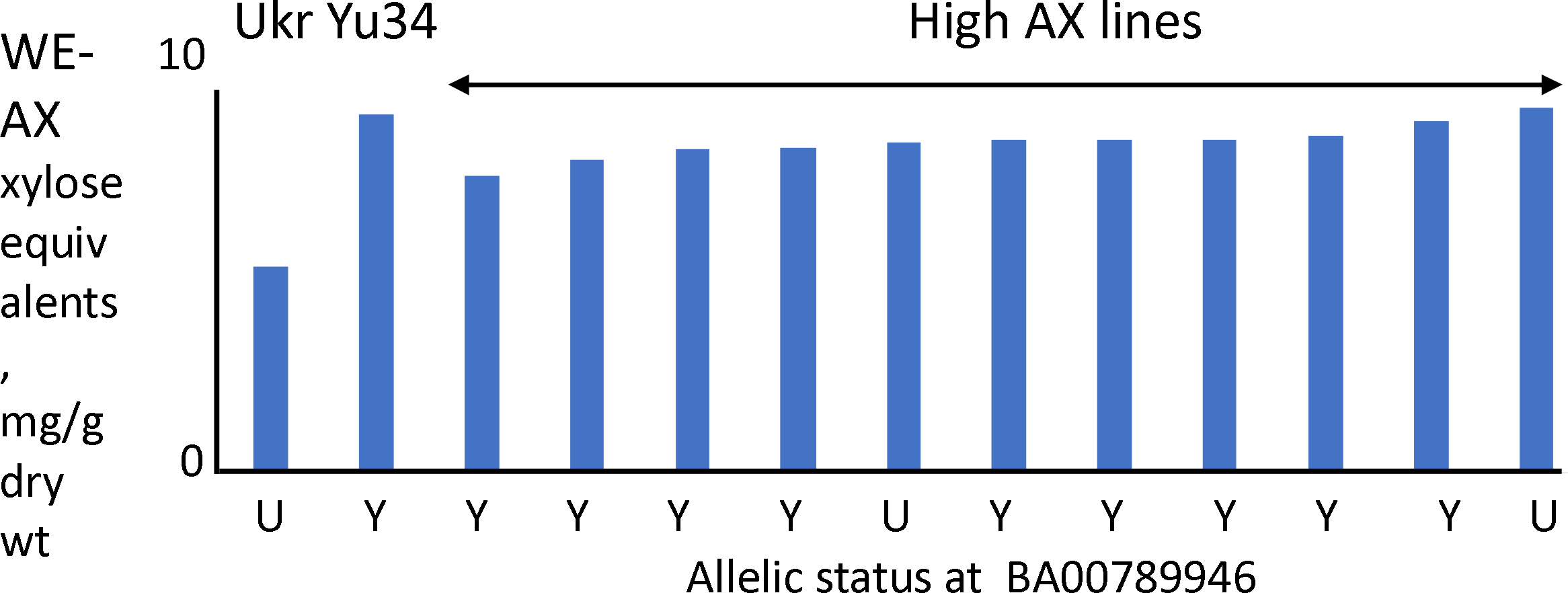
Contents of WE-AX (determined as pentosane expressed as xylose equivalents) and GC-FID in white flours of 11 high AX lines derived from the cross Yumai 34 x Ukrainka using biochemical analysis for selection. Data are the means of samples grown at Martonvasar over 3 years (2013-2015) as previouly reported by Tremmel-Bede et al. (2017).

## Discussion

We have used four crosses with the high fibre wheat cultivar Yumai 34 to identify QTL for high RV and total AX fibre in white flour. Although a number of QTL were mapped, which is consistent with earlier studies, most of these were only detected in one or two crosses. However, all four crosses showed strong QTLs for high AX/RV on chromosome 1B, with Yumai 34 being the increasing parent, although this mapped to the 1BS in the Y34Ukr and to 1BL in the other crosses. Furthermore, a KASP marker for the high AX Yumai 34 allele was validated by analysis of high AX lines derived from Yumai 34 but selected by conventional biochemical analysis.

Previous studies have also shown QTLs for RV on chromosome 1B, including a QTL on 1BL with a heritability of 27.4% in one population (Quraishi et al., 2011). This may correspond to the 1BL QTL in Yumai 34, with the Yumai allele giving higher levels of AX and RV. The two QTL on 1B described here appear to be genuinely independent. We investigated whether a translocation present in Ukrainka might cause 1BL markers in Y34Ukr to link with markers on 1BS, but there was no evidence for this (data not shown). Moreover, the BA00789946 KASP assay is assigned to 1BL when mapped using the Y34Ukr population, as in the other three crosses. Nevertheless, this marker has strong predictive power for high fibre in the progeny selected from crosses between Yumai 34 and Ukrainka, which supports the QTL for RV on 1BL in this population even though the LOD score was below the cut-off.

The most likely explanation for the identification of a TOT-AX QTL on 1BS in the cross with Ukrainka is that Yumai 34 does indeed carry AX increasing effects at both ends of chromosome 1B. However, detecting linked QTLs requires higher statistical power and it is likely that we did not detect the TOT-AX QTL on 1BL as significant in the same cross due to insufficient power.

It is notable that only one cross, between Yumai 34 and the high AX cultivar Valoris, showed significant transgressive segregation. Previous analysis of a cross between Valoris and Isengrain (which has a normal AX level) identified a QTL on 6BL which explained 58.4% of the heritability (Charmet et al., 2009; Quraishi et al., 2011). This may correspond to the QTL identified on 6B in the Yumai 34 × Valoris cross, although this QTL was mapped to 6BS not 6BL. The presence of the high AX allele of the 1B QTL in Valoris probably accounted for the presence of transgressive segregation, which indicates that it should be possible to stack the 1B and 6B QTLs, and possibly also other high AX QTLs.

## Experimental procedures

### Production and growth of materials

Populations of Yumai 34 × Ukrainka, Yumai 34 × Claire and Yumai 34 × Valoris were grown on three sites in the UK from 2012-2015 at Rothamsted Research, Hertfordshire, AL5, 2JQ), Church Farm, Norfolk (NR9 3PY), UK, and KWS UK Ltd, Thriplow, Cambridgeshire, SG8, 7RE, UK. At each field trial site three replicate 1m^2^ plots were grown per line in a randomised plot design. The populations Yumai 34 × Valoris and Yumai 34 × Altigo were also grown at INRA, Clermont Ferrand, France in two replicates of 7.5m^2^ plots, 2013.

The cultivars Yumai 34, Hereward, Manital and Soisson varieties were also grown in 1.8 × 1m^2^ plots at Rothamsted Research in 2011/12. Analyses of non-starch polysaccharides in white flours of these samples are given in Table 1 and representative enzymatic fingerprinting data in Supplementary Table S1.

High fibre lines selected from a cross between Yumai 34 and Ukrainka were grown at the Centre for Agricultural Research, Martonvásár.

### Milling

White flour was produced using a Chopin CD1 mill. Grains were brought to room temperature and moisture content determined using Bruker Minispec mq-20 NMR analyser using an in-house developed calibration. 50g of grain was conditioned to 16.5% moisture overnight prior milling. First break and first reduction flours were combined to give the white flour fraction.

Wholemeal flour was produced from mature grain of the crosses Yumai 34 × Ukrainka, Yumai 34 × Claire using a Retch centrifugal mill with a 250μm sieve. Wholemeal flours of the crosses Yumai 34 × Altigo and Yumai 34 × Valoris were produced by ball milling at 8°C with a 500μm sieve.

All wholemeal and white flour samples were immediately stored at −20°C prior to use.

### Determination of relative viscosity (RV)

RV of aqueous extracts of white flours and of wholemeal flours of the crosses Yumai 34 × Ukrainka and Yumai 34 × Claire were determined at 30 °C using an automated viscometer (AVS 370, SI Analytics, Germany) fitted with a Micro Ostwald capillary (2 mL, 0.43 mm) and WinVisco software (Freeman et al., 2016). Relative viscosities of wholemeal flours of the crosses Yumai 34 × Altigo and Yumai 34 × Valoris were determined at 30 °C using an automated viscometer (AVS 310, CV (%) 23 22 8 Schott Gerate, Germany) fitted with an Ostwald capillary (2 mL, 0.4 mm) (Saulnier et al., 1995).

### Arabinoxylan analysis

Total and water-un extractable AX in white flour were determined by monosaccharide analysis, by GC-FID of alditol acetates as described by Gebruers et al. (2009) using the method of Englyst et al. (1995). Results were adjusted for the presence of arabinogalactan as described in Gebruers et al. (2009).

Total and water-extractable (WE) AX were also determined in wholemeal and white flour samples from 3 replicate plots with two technical replications using the method described in Finnie et al., 2006 based upon the method of Douglas (1981).

Enzymatic fingerprinting of AX was as described by Lovegrove et al. (2013). White flour was digested using a mixture of endoxylanase and lichenase (β-glucanase) to release arabinoxylan oligosaccharides (AXOS) and glucan fragments comprising 3 and 4 residues (G3, G4), respectively. These were separated using a Carbopac PA-1 (Dionex) column with dimensions 2 mm × 250 mm and the flow rate of 0.25 mL/min based upon the original method of Ordaz-Ortiz et al. (2005). At least two technical replicates of each biological replicate were analysed. The areas under the AXOS peaks were combined to determine TOT-AX (expressed in arbitrary units).

### Markers and mapping

Genetic maps were constructed using KASP single nucleotide polymorphism (SNP) markers or the Axiom 35K breeders array. New KASP markers were developed, where necessary, from SNPs detected by Axiom markers using assays shown on Cerealsdb predesigned using polymarker (Ramirez et al 2015). The KASP primers used for SNP BA00789946 (shown as AX-94524314 on the genetic maps presented here) were: acaactaactatgcaagtgcca (FAM labelled), acaactaactatgcaagtgccg (VIC labelled) and ggatgacacatctcaagaaaaagaa. Standard conditions for the KASP PCR were used, with a hydrocycler and a touchdown of 65°C to 57°C.

## Supporting information

Additional Supporting Information may be found online in the supporting information tab for this article:

**Table S1**. Relative amounts of AXOS and GOS released by enzyme digestion of white flour from four wheat cultivars

**Figure S1**. Correlation Matrix of 10 lines from the Yumai x Ukrainka population (2013/2014) grown at Rothamsted Research of relative viscosity, water extractable (WE) and total AX measured colorometrically (pentosan assay), and total AX by enzyme fingerprinting in wholemeal and white flour.

**Figure S2**. Relative viscosity of RILs from the crosses Yumai 34 x Claire (top) and Yumai 34 x Ukrainka

## Authors’ contributions

Alison Lovegrove: Peter R Shewry, Simon Griffiths: designed experiments, supervised analyses, wrote paper

Luzie Wingen: analysed mapping data

Amy Plummer, Abigail Wood, Diana Passmore, Ondrej Kosik, Jackie Freeman, Mehmet Ulker, Till Pellny: carried out field trials and biochemical analyses of grain

Karolina Tremmel-Bede, Marianna Rakszegi, Zoltán Bedő: constructed Y34×U mapping population

Marie-Reine Petterant, Gilles Charmet: constructed Y34Alt and Y34Val mapping populations, carried out field trials and determined RV.

Michelle Leverington Waite, Simon Orford: constructed Y34Cl mapping population; carried out field trials, prepared samples of all populations for marker analysis.

Rowan Mitchell: designed and supervised experiments

Amanda Burridge: carried out marker analysis of mapping populations

## Acknowledgements

Rothamsted Research and the John Innes Centre receive grant-aided support from the Biotechnology and Biological Sciences Research Council (BBSRC) of the UK and the work reported here forms part of the Designing Future Wheat Institute Strategic Programme [BB/P016855/1], with additional support from Innovate UK grant, L005654/1 “High fibre wheat for healthier white bread”. We are grateful to Dr.Jacob Lage and colleagues at KWS UK Ltd for field trials of populations.

## References

Anderson JW, Baird P, Davis RH Jr, Ferreri S, Knudtson M, Koraym A, Waters V, Williams CL (2009) Health benefits of dietary fiber. Nutrition Reviews 67: 188–205.

Andersson AAM, Andersson R, Piironen V, Lampi A-M, Nyström L, Boros D, Fraś A, Gebruers K, Courtin C, Delcour J, Rakszegi M, Bedő Z, Ward JL, Shewry P, Åman P. (2012) Contents of dietary fibre components and their relation to associated bioactive components in whole grain wheat samples from the HEALTHGRAIN diversity screen. Food Chemistry 136: 1243–1248.

Appels R, Eversole K, Feuillet C, Keller B, Rogers J, Stein N, Pozniak CJ, Choulet F, Distelfeld A, Poland J, Ronen G. (2018) Shifting the limits in wheat research and breeding using a fully annotated reference genome. Science 361: (6403) pii.eaar7191.

Aune D, Chan DSM, Lau R, Vieira R, Greenwood DC, Kampman E, Norat T. (2011) Dietary fibre, whole grains, and risk of colorectal cancer: systematic review and dose-response meta-analysis of prospective studies. British Medical Journal 343: d6617 doi: 10.1136/bmj.d6617.

Aune D, Keum N, Giovannucci, Fadnes LT, Boffetta P, Greenwood DC, Tonstad S, Vatten LJ, Riboli E, Norat T. (2016) Whole grain consumption and risk of cardiovascular disease, cancer, and all cause and cause specific mortality: systematic review and dose-response meta-analysis of prospective studies British Medical Journal. 353: i2716. doi: 10.1136/bmj.i2716.

Bates B, Lennox A, Prentice A, Bates C, Page P, Nicholson S, Milne A. Swan G. (2014). National diet and nutrition survey rolling programme (NDNS RP). Results from years 1–4 (combined) for Scotland (2008/9-2011/12). Public Health England and Food Standards Agency in Scotland.

Cade JE, Burley VJ, Greenwood DC and U.K. Women’s Cohort Study Steering Group. (2007). Dietary fibre and risk of breast cancer in the UK Women’s Cohort Study. International Journal of Epidemiology 36: 431–438

Charmet G, Masood-Quraishi U, Ravel C, Romeuf I, Balfourier F, Perretant MR, Joseph JL, Rakszegi M, Guillon F, Sado PE, Bedo Z, Saulnier L (2009). Genetics of dietary fibre in bread wheat. Euphytica 170: 155–168.

Cooper DN, Martin R, Keim L. (2015) Does whole grain consumption alter gut microbiota and satiety? Healthcare 3: 364–92.

Douglas SG. 1981. A rapid method for the determination of pentosans in wheat flour.Food Chemistry 7: 139–145.

Englyst HN, Quigley ME, Hudson GJ. (1994). Determination of dietary fibre as non-starch polysaccharides with gas-liquid chromatographic, high-performance liquid chromatographic or spectrophotometric measurement of constituent sugars.Analyst 119:1497–1509.

Finnie SM., Bettge AD, Morris CF. (2006). Influence of cultivar and environment on water-soluble and water-insoluble arabinoxylans in soft wheat. Cereal Chemistry 83: 617–623.

Freeman J, Lovegrove A, Wilkinso MD, Saulnier L, Shewry PR, Mitchell RA. (2016). Effect of suppression of arabinoxylan synthetic genes in wheat endosperm on chain length of arabinoxylan and extract viscosity. Plant Biotechnology Journal 14:109–116.

Gebruers K, Dornez E, Boros D, Fraś A, Dynkowska W, Bedő Z, Rakszegi M, Delcour JA Courtin CM. (2008) Variation in the content of dietary fiber and components thereof in wheats in the HEALTHGRAIN diversity screen. Journal of Agricultural and Food Chemistry 56: 9740–9749.

Gebruers K, Courtin CM Delcour, JA. (2009). Quantitation of arabinoxylan and their degree of branching using gas chromatography. In: Analysis of Bioactive Components in Small grain Cereals. (Shewry PR and Ward JL. eds), pp177–190. St Paul: AACC International.Inc.

Hajishafiee M, Saneei P, Benisi-Kohansal S, Esmaillzadeh A. (2016) Cereal fibre intake and risk of mortality from all causes, CVD, cancer and inflammatory diseases: a systematic review and meta-analysis of prospective cohort studies. British Journal of Nutrition 116: 343—352.

Helldán A, Raulio S, Kosola M, Tapanainen H, Ovaskainen M.-L, Virtanen S. (2012) The National FINDIET 2012 Survey. National Institutes of Health and Welfare, Helsinki, Finland.

International Wheat Genome Sequencing Consortium. (2014) A chromosome-based draft sequence of the hexaploid bread wheat (Triticum aestivum) genome. Science 345: 1251788. doi: 10.1126

Laperche A, Brancourt-Hulmel M, Heumez E, Gardet O, Hanocq E, Devienne-Barret F, and Le Gouis J. (2007). Using genotype x nitrogen interaction variables to evaluate the QTL involved in wheat tolerance to nitrogen constraints. Theoretical and Applied Genetics 115: 399–415.

Lovegrove A, Wilkinson MD, Freeman J, Pellny TK, Tosi P, Saulnier L, Shewry PR, Mitchell RAC. (2013) RNA interference suppression of genes in glycosyl transferase families 43 and 47 in wheat starchy endosperm causes large decreases in arabinoxylan content. Plant Physiology 163: 95—107.

Marcotuli I, Houston K, Waugh R, Fincher GB, Burton RA, Blanco A, and Gadaleta A. (2015). Genome wide association mapping for arabinoxylan content in a collection of tetraploid wheats. PLOS ONE DOI: 10.1371/journal.pone.0132787

Mares DJ, Stone BA. (1973) Studies on wheat endosperm. I. Chemical composition and ultrastructure of the cell walls. Australian Journal of Biological Science 26: 793–812.

Martinant JP, Billot A, Bouguennec A, Charmet G, Saulnier L, Branlard G. (1999) Genetic and environmental variations in water-extractable arabinoxylans content and flour extract viscosity. J Cereal Science 30: 45–48.

Mattei J, Malik V, Wedick NM, Hu FB, Spiegelman D, Willett WC, Campos H, and Global Nutrition Epidemiologic Transition Initiative. (2015) Reducing the global burden of type 2 diabetes by improving the quality of staple foods: The Global Nutrition and Epidemiologic Transition Initiative. Globalization and Health 11: 23 DOI 10.1186/s12992-015-0109-9

Nyugen V-L, Huynh B-L, Wallwork H, Stangoulis J. (2011) Identification of quantitative trait loci for grain arabinoxylan concentration in bread wheat. Crop Science 51: 1143–1150.

Ordaz-Ortiz JJ, Devaux MF, Saulnier L. (2005) Classification of wheat varieties based on structural features of arabinoxylans as revealed by endoxylanase treatment of flour and grain. Journal of Agricultural and Food Chemistry 53: 8349—8356

Ordaz-Ortiz JJ, Saulnier L. (2005) Structural variability of arabinoxylans from wheat flour. Comparison of water-extractable and xylanase-extractable arabinoxylans. Journal of Cereal Science 42:119–125.

Perretant MR, Cadalen T, Charmet G, Sourdille P, Nicolas P, Boeuf C, Tixier MH, Branlard G, Bernard S, Bernard M. (2000). QTL analysis of bread-making quality in wheat using a doubled haploid population. Theoretical and Applied Genetics 100:1167–1175.

Quraishi U-M, Abrouk M, Bolot S, Pont C, Throude M, Guilhot N, Confolent C, Bortolini F, Praud S, Murigneux A, Charmet G and Salse J (2009), ‘Genomics in cereals: from genome-wide conserved orthologous set (COS) sequences to candidate genes for trait dissection’, Functional and Integrated Genomics, 9: 473–484.

Quraishi U-M, Murat F, Abrouk M, Pont C, Confolent C, Oury F X, Ward J, Boros D, Gebruers K, Delcour J A, Courtin C M, Bedo Z, Saulnier L, Guillon F, Balzergue S, Shewry P R, Feuillet C, Charmet G and Salse J (2010). ‘Combined meta-genomics analyses unravel candidate genes for the grain dietary fibre content in bread wheat (Triticum aestivum L.). Functional and Integrated Genomics, 10: doi 1007/s10142-010-0183-2.

Ramirez-Gonzalez RH, Uauy C Caccamo, M. PolyMarker: a fast polyploid primer design pipeline (2015) Bioinformatics 31: 2038–2039.

Reynolds A, Mann J, Cummings J, Winter N, Mete E, Morenga L T (2019). Carbohydrate quality and human health: a series of systematic reviews and meta-analyses. Lancet, doi.org/10.1016/S0140-6736(18)31808–9.

Shewry PR (2013). Improving the content and composition of dietary fibre in wheat. In: Fibre-Rich and Wholegrain Foods: Improving Quality (Delcour JA, Poutanen K. eds), pp. 153–169. Oxford: Woodhead Publishing,

Steer, Thane C, Stephen A, Jebb S. (2008) Bread in the diet: consumption and contribution to nutrient intakes of British adults. Proceedings of the Nutrition Society 67: E363.

Saulnier L, Peneau N Thibault J-F. (1995). Variability in grain extract viscosity and water-soluble arabinoxylan content in wheat. Journal of Cereal Science 22: 259–264.

Tremmel-Bede K, Lang L, Torok K, Tomoskosi S, Vida G, Shewry PR, Bedo, Z, Rakszegi M. (2017). Development and characterization of wheat lines with increased levels of arabinoxylan. Euphytica 213: 291.

Yang L, Zhao D, Yan J, Zhang Y, Xia X, Tian Y, He Z, Zhang Y. (2015). QTL mapping of grain arabinoxylan contents in common wheat using a recombinant inbred line population. Euphytica DOI 10.1007/s10681-015-1576-z

Ye EQ, Chacko SA, Chou EL, Kugizaki M, Liu S. (2012). Greater whole-grain intake is associated with lower risk of type 2 diabetes, cardiovascular disease, and weight gain. Journal of Nutrition 142:1304—1313

Wolever TMS, Tosh SM, Gibbs AL, Brand-Miller J, Duncan AM, Hart V, Lamarche B, Thomson BA, Duss R, Woo P.J. (2010). Physiochemical properties of oat β-glucan influence its ability to reduce serum LDL cholesterol in humans: a randomized clinical trial. American Journal of Clinical Nutrition 92: 723–732.

Wood PJ. (2007) Cereal β-glucans in diet and health. Journal of Cereal Science 46: 230’238

